# Circadian clock regulates epidermal endocrine system in homeostatic skin pigmentation

**DOI:** 10.1101/2025.11.20.689605

**Authors:** Anya Zhu, Lingli Yang, Sylvia Lai, Fei Yang, Daisuke Tsuruta, Ichiro Katayama

**Author notes:** Correspondence to: L Yang, Department of Pigmentation Research and Therapeutics, Graduate School of Medicine, Osaka Metropolitan University, Osaka 5450051, Japan.

## Abstract

The circadian clock regulates various physiological processes in the skin, including local hormone synthesis and pigment production. However, the interplay between circadian regulation and epidermal endocrine signaling remains poorly understood. In this study, we investigated the functional roles of two core circadian transcriptional regulators, Basic helix-loop-helix ARNT-like protein 1 (BMAL1) and CLOCK circadian regulator (CLOCK), in human epidermal homeostasis. Primary normal human epidermal melanocytes were cultured and analyzed using quantitative real-time PCR, Western blotting, and immunofluorescence. Both BMAL1 and CLOCK were abundantly expressed in melanocytes. siRNA-mediated knockdown of either gene led to a significant reduction in melanin synthesis and altered the expression of hormone-related factors, notably decreasing the expression of vitamin D receptor (VDR) and its associated pathways. These molecular alterations were closely correlated with changes in melanogenic activity. Consistently, immunohistochemical analysis of vitiligo patient skin revealed reduced expression of the vitamin D-related enzyme CYP27A1 in melanocytes. Collectively, our findings demonstrate that circadian clock disruption impairs pigment production partly through modulation of vitamin D signaling, uncovering a previously unrecognized link between the cutaneous circadian system and endocrine regulation of melanogenesis.

## 1. Introduction

All living organisms exhibit circadian rhythms that regulate a wide range of physiological and behavioral processes^1^. At the cellular level, the circadian clock consists of transcriptional-translational feedback loops that generate rhythmic gene expression^2^. The core regulatory loop involves BMAL1 and CLOCK, which form a heterodimer to activate the transcription of Period (PER1-3) and Cryptochrome (CRY1-2) genes. The translated PER and CRY proteins heterodimerize and translocate into the nucleus, where they inhibit their own transcription by interacting with the CLOCK: BMAL1 complex. In addition, the CLOCK:BMAL1 complex induces RORα and REV-ERBα, which positively and negatively regulate BMAL1 transcription, respectively, forming an auxiliary loop that stabilizes circadian oscillation^3^. This intrinsic system enables cells to anticipate and adapt to daily environmental changes such as light, temperature, and nutrient availability.

Beyond the central pacemaker in the suprachiasmatic nucleus (SCN), peripheral circadian oscillators have been identified in various organs, including the skin, liver, and adrenal gland^4,5^. In the skin, keratinocytes, melanocytes, and fibroblasts display autonomous circadian rhythmicity that regulates cellular proliferation, DNA repair, differentiation, and pigmentation^6–11^. Several physiological processes of the skin, such as transepidermal water loss, barrier recovery, and melanin synthesis, exhibit diurnal variation, highlighting the importance of peripheral clocks in maintaining cutaneous homeostasis^12–15^. The rhythmic expression of clock genes in the epidermis not only reflects systemic cues but also responds directly to local stimuli such as ultraviolet (UV) exposure and temperature fluctuations, linking environmental light cycles to molecular processes in the skin^14,16–18^.

Accumulating evidence indicates that circadian disruption contributes to various skin disorders^15,19–21^. Epidemiological studies have reported that night-shift workers have an increased incidence of psoriasis^22^. Moreover, polymorphisms in BMAL1 have been identified in patients with non-segmental vitiligo, suggesting that dysregulation of circadian genes may predispose individuals to pigmentary or autoimmune skin diseases^23^.

In addition to its barrier and sensory roles, the skin also functions as a highly active local endocrine organ^24–27^. It is capable of synthesizing, metabolizing, and responding to a variety of hormones and hormone-like molecules, including vitamin D₃, thyroid hormones, melatonin, cortisol, estrogens, and light-sensitive opsins. These endocrine factors, along with their biosynthetic enzymes and receptors, are expressed in multiple skin cell types, particularly keratinocytes and melanocytes. They modulate pigmentation, cellular proliferation, oxidative stress, and immune responses, thereby influencing both physiological adaptation and pathological processes^26,28–31^. Notably, many of these hormone systems are under circadian regulation in other tissues, suggesting that the cutaneous endocrine microenvironment may also be governed by local circadian control^24–29^. However, the molecular interplay between core circadian regulators, melanocyte function, and local endocrine signaling remains poorly understood.

Melanocytes are the pigment-producing cells in the epidermis. Recent studies have shown a specific peripheral clock characterized by rhythmic expression of core clock genes like BMAL1, PER1, and PER2, and that silencing these genes stimulates melanogenic activity, linking clock genes to melanin synthesis regulation^7,31,32^. Nevertheless, it remains unclear whether disruption of circadian clock genes affects the biosynthesis or receptor signaling of local hormones, such as vitamin D₃, thyroid hormones, melatonin, cortisol, estrogens, and opsins, in melanocytes.

In this study, we investigated how disruption of BMAL1 and CLOCK affects human melanocyte function and the biosynthesis and signaling of various local hormones, including vitamin D₃, thyroid hormone, melatonin, cortisol, estrogen, and opsins. This work aims to elucidate how circadian clock genes integrate endocrine pathways to regulate pigmentation and epidermal homeostasis.

## 2. Materials and Methods

### 2.1. Cell lines and cell culture

Human foreskin keratinocytes immortalized by infection with the pSV40 ori (PSVK1 cells) were purchased from the Japanese Collection of Research Bioresources (JCRB, Osaka, Japan) and cultured in KBM-Gold Keratinocyte Basal Medium (Lonza, Walkersville, MD, USA) supplemented with the KGM-Gold Bullet Kit (Lonza). HEKn (normal neonatal human epidermal keratinocytes) and HEKa (normal adult human epidermal keratinocytes) cells were obtained from Cascade Biologics^TM^ (Portland, OR, USA) and cultured in Epilife medium supplemented with Human Keratinocyte Growth Supplement (HKGS) (GIBCO, Gaithersburg, MD, USA). The ISO-HAS-B (ISO1) skin angiosarcoma cell line was obtained from the Cell Resource Center for Biomedical Research of the Cell Bank (Sendai, Japan) and cultured in high-glucose Dulbecco’s modified Eagle’s medium (DMEM) supplemented with 10% fetal bovine serum and 1% penicillin–streptomycin (Thermo Fisher). NHDF cells (human normal dermal fibroblasts) were acquired from Invitrogen (Thermo Fisher Scientific, Carlsbad, CA, USA) and cultured in low-glucose DMEM supplemented with 10% fetal bovine serum and 1% penicillin-streptomycin (Thermo Fisher, Waltham, MA, USA). HKA (cell line derived from a keratoacanthoma of human skin), Mewo, and G361 (human melanoma cell lines) cells were procured from the Japanese Collection of Research Bioresources (JCRB, Osaka, Japan) and maintained in high-glucose DMEM supplemented with 10% fetal bovine serum and 1% penicillin-streptomycin (Thermo Fisher, Waltham, MA, USA). HaCaT cells (human immortalized keratinocytes) were obtained from the American Type Culture Collection (ATCC, Manassas, VA, USA). HEMn-MPs (normal human primary skin epidermal melanocytes from moderately pigmented neonatal foreskin) were obtained from Invitrogen (Thermo Fisher Scientific, Carlsbad, CA, USA) and cultured in Medium 254 (M-254-500; Thermo Fisher Scientific) supplemented with 1% (*v/v*) human melanocyte growth supplement (Thermo Fisher Scientific). HMVECs (human microvascular endothelial cells) were procured from Lonza (Walkersville, MD, USA) and cultured in endothelial basal medium-2 (EBM-2, Lonza). Detailed information about the sources of all cultured cells, including age and generation, is provided in Figure 1F. All cultures were maintained at 37°C in a CO_2_ incubator with an atmosphere of 5% CO_2_.

**Figure 1.**
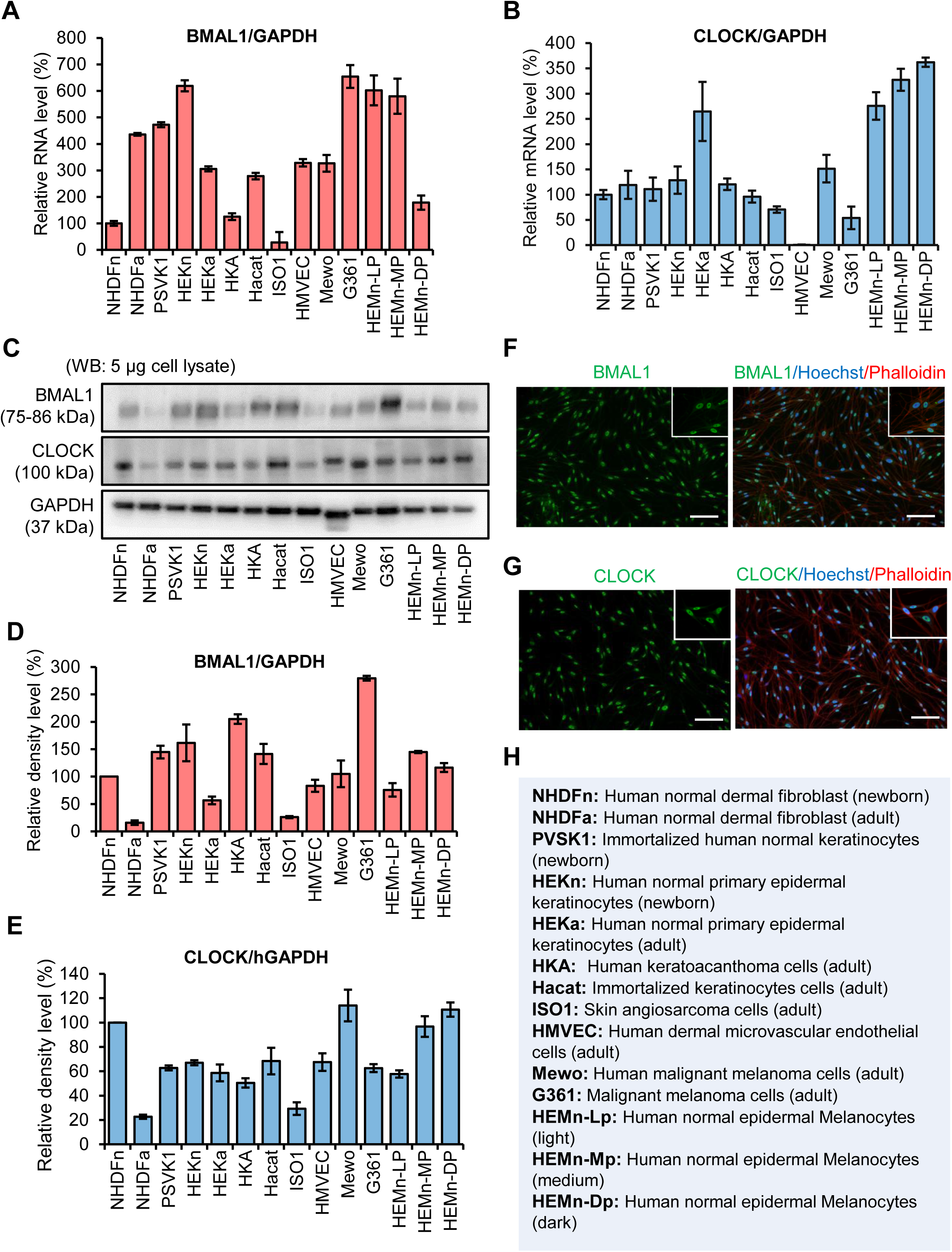
Expression of BMAL1 and CLOCK across diverse human skin cell types. Quantitative real-time PCR analysis of BMAL1 **(A)** and CLOCK **(B)** mRNA expression in fourteen human skin-derived cell types. Transcript levels were normalized to GAPDH and presented relative to NHDFn. Representative Western blot analysis showing BMAL1 and CLOCK protein expression in whole-cell lysates from the indicated cultured cells, with GAPDH serving as the loading control **(C)**. Densitometric quantification of BMAL1 **(D)** and CLOCK **(E)** protein levels normalized to GAPDH. Immunofluorescence staining of cultured melanocytes (HEMn-LP) showing the intracellular localization of BMAL1 **(F)** and CLOCK **(G)**. BMAL1/CLOCK (green), Hoechst 33342 (blue), and Phalloidin (red). Insets highlight representative cells at higher magnification. Scale bar: 100 μm. Summary table listing detailed information on all cell lines used in this study, including species and tissue origin. Data in **A**, **B**, **D**, and **E** are shown as mean ± SD.

### 2.2. Human Skin Specimens

Paraffin-embedded tissue sections of lesional skin from confirmed vitiligo patients in the progressive state (n = 4) and samples from corresponding sites of healthy donors (n = 4) were used in histo-immunofluorescence staining. Written informed consent was obtained from all participants prior to study inclusion. The study was approved by the ethics committee of the Osaka Metropolitan University Faculty of Medicine (No. 4152).

### 2.3. RNA interference

For siRNA-mediated knockdown of BMAL1 and CLOCK, HEMn-MPs were transfected with 30 nM pre-designed Silencer^®^ Select siRNAs (si-BMAL1: HSS100702; si-CLOCK: HSS114292; Thermo Fisher Scientific) using Lipofectamine® RNAi MAX (Invitrogen, Thermo Fisher Scientific) according to the manufacturer’s instructions.

### 2.4. RNA isolation and real-time RT-PCR analysis

Total RNA was extracted from cell pellets using the Maxwell^®^ 16 LEV simplyRNA Tissue Kit (Promega, Madison, WI, USA), following the manufacturer’s protocol. RNA integrity was confirmed through gel electrophoresis. Subsequently, 100 ng of total RNA was reverse-transcribed into first-strand cDNA using the ReverTra Ace® qPCR RT Master Mix (TOYOBO, Osaka, Japan).

The primers used for real-time PCR were as follows: human BMAL1 sense, 5’-GCTTCTGCACAATCCACAGC-3’ and antisense, 5’-CACCCTGATTTCCCCGTTCA-3’; human CLOCK sense, 5’-CGAGCGCTCCCGAATTTTTA-3’ and antisense, 5’-AGGTATCTAGTGAGACTTGCCA-3’; human short-wave-sensitive opsin 1 (OPN1SW) sense, 5’-TATCTCTTCAGTGGGGCCGT-3’ and antisense, 5’-GGCTGCCGCAACTTTTTGTA-3’; human opsin-2 (OPN2) sense, 5’-CATGACCATCCCAGCGTTCT-3’ and antisense, 5’-CTTGGACACGGTAGCAGAGG-3’; human opsin-3 (OPN3) sense, 5’-CTACAAGTTCCAGCGGCTCC-3’ and antisense, 5’-CGAAGGTAAAGGTGACCCCG-3’; human peropsin sense, 5’-GATACGCAGGCTGTCAGGTT-3’ and antisense, 5’-GGCAGATGGTCAGGTATCGG-3’; human thyroid hormone receptor alpha (TRα) sense, 5’-GGAGAAGGGTGACGTTGGAA-3’ and antisense, 5’-TTTCATCCTTGTGGGGGTTCA-3’; human thyroid hormone receptor beta (TRβ) sense, 5’-GCGATTTCCTTCTGGTTGGC-3’ and antisense, 5’-AGTGCGGTTTCCTTATGGCT-3’; human tryptophan hydroxylase 1 (TPH1) sense, 5’-TCTACCCAACCCATGCTTGC-3’ and antisense, 5’-AAGTAACCAGCCACAGGACG-3’; human tryptophan hydroxylase 2 (TPH2) sense, 5’-TTGGGGTGTTGTATTCCGGG-3’ and antisense, 5’-CCGTGAAGCCAGACCTTTCT-3’; human Hydroxyindole-O-methyltransferase (HIOMT) sense, 5’-ACGGCTGGATTGGAGACAAG-3’ and antisense, 5’-CTCGGCGAGAAGGTCAAACA-3’; human melatonin receptor 1a (MTNR1A) sense, 5’-CACCATCGTGGTGGACATCCT-3’ and antisense, 5’-GCACCAACGGGTACGGATA-3’; human melatonin receptor 1b (MTNR1B) sense, 5’-GCTGCCCAACTTCTTTGTGG-3’ and antisense, 5’-GACACGACAGCGATAGGGAG-3’; human cytochrome P450 family 27 subfamily A member 1 (CYP27A1) sense, 5’-CCTTCGTCAGATCCATCGGG-3’ and antisense, 5’-GGGCCTCCATATCTTCGAGC-3’; human cytochrome P450 family 27 subfamily B member 1 (CYP27B1) sense, 5’-CCTGACCCACTTCCTGTTCC-3’ and antisense, 5’-CTGAGTGGAGTGCTGTCTGG-3’; human cytochrome P450 family 24 subfamily A member 1 (CYP24A1) sense, 5’-TGCCAGCGATAATACGCCTC-3’ and antisense, 5’-TCCCAGGCCATTCTAAGCAC-3’; human vitamin D receptor (VDR) sense, 5’-CGCCCACCATAAGACCTACG-3’ and antisense, 5’-GGGAGTGTGTCTGGAGTTGG-3’; human nuclear receptor subfamily 3, group C, member 1 (NR3C1) sense, 5’-GAGGGAAGGAAACTCCAGCC-3’ and antisense, 5’-TCAGCTAACATCTCGGGGAA-3’; human nuclear receptor subfamily 3, group C, member 2 (NR3C2) sense, 5’-TGCAAAAGAACCCTCGGTCA-3’ and antisense, 5’-GCGTGGAGAGCAGATTTTCG-3’; human G protein-coupled estrogen receptor 1 (GPER1) sense, 5’-GGGACAACTGCGGTGATGAT-3’ and antisense, 5’-GGATCCGCACATGACAGGTT-3’; human Estrogen receptor alpha (ERα) sense, 5’-AACAGGCTCGAAAGGTCCAT-3’ and antisense, 5’-CAGTCCCGGAGAATGTGAAGA-3’; human Estrogen receptor beta (ERβ) sense, 5’-CGTGACCGATGCTTTGGTTT-3’ and antisense, 5’-AGCAGATGTTCCATGCCCTT-3’; human Microphthalmia-associated Transposition factor (MITF) sense, 5’-CCGGGCTCTGTTCTCACTT-3’ and antisense, 5’-CATGAAACTCCTCCCCGACT-3’; human Dopachrome Tautomerase (DCT) sense, 5’-CCCATTTTTGTGGTTCTTCATTCC-3’ and antisense, 5’-CGATTGTGACCAATAGGGGC-3’; human Tyrosinase (TYR) sense, 5’-TGACTCCAATTAGCCAGTTCCT-3’ and antisense, 5’-GACAGCATTCCTTCTCCATCAG-3’; human Tyrosinase-related protein (TYRP1) sense, 5’-CTCAATGGCGAGTGGTCTGT-3’ and antisense, 5’-TTCCAAGCACTGAGCGACAT-3’; human Premelanosome Protein 17 (Pmel17) sense, 5’-CTATGTGCCTCTTGCTCATTCC-3’ and antisense, 5’-TGCTTGTTCCCTCCATCCA-3’; human Stem cell factor receptor (c-Kit) sense, 5’-GCACAATGGCACGGTTGAAT-3’ and antisense, 5’-GGTGTGGGGATGGATTTGCT-3’; human Endothelin receptor type B (EDNRB) sense, 5’-CTAGGCTCTGAAACTGCGGC-3’ and antisense, 5’-GGCGTCATTATCTCTGCGGT-3’; and human glyceraldehyde 3-phosphate dehydrogenase (GAPDH) sense, 5’-GACAGTCAGCCGCATCTTCT-3’ and antisense, 5’-GCGCCCAATACGACCAAATC-3’. Real-time PCR was performed using a QuantStudio^®^5 Real-time PCR System (Applied Biosystems, CA, USA). Each reaction was conducted in triplicate in three independent experiments. The housekeeping gene, GAPDH, was employed as an internal control for normalization.

### 2.5. Western blot analysis

For protein sample preparation, cell pellets were extracted as described previously^33^, and 5 μg of the extracted protein was utilized for Western blot analysis. Primary antibodies were employed at the following dilutions: anti-BMAL1 (#14020; CST, Danvers, MA, USA) at 1:500, anti-CLOCK (#5157S; CST) at 1:500, anti-Pmel17 (sc377325, Santa Cruz Biotechnology, Texas, USA) at 1:500, anti-TYRP1 (sc58438, Santa Cruz Biotechnology) at 1:500, anti-Melan-A (ab51061, Abcam, Cambridge, UK), anti-CYP27A1 (ab126785; Abcam) at 1:500, and anti-GAPDH (#2118; CST) at 1:1000. The anti-GAPDH antibody was employed as a loading control.

### 2.6. Immunofluorescence staining of cells

HEMn-MPs were seeded on six-well plates and allowed to grow to confluence. After 24 hours of incubation (or 48 hours following siRNA transfection), the cells underwent a thorough washing with ice-cold PBS and were then fixed with 4% paraformaldehyde for 5 minutes at room temperature. Fixed cells were then incubated with primary antibodies against BMAL1 (#14020; CST) or CLOCK (#5157S; CST) at a dilution of 1:500. Actin filaments were labeled using Alexa Fluor 555 Phalloidin (A34055, Invitrogen; 1:100 dilution), and nuclei were counterstained with Hoechst 33342 (Invitrogen; 1:500 dilution). Fluorescence images were acquired using a Biozero BZ-8100 confocal microscope (Keyence Corporation, Osaka, Japan).

### 2.7. 3-[4-dimethylthiazol-2-yl]-2,5-diphenyltetrazolium bromide (MTT) assay

HEMn-MP cells (2.5 × 10^4^ cells/well) were cultured in 96-well flat-bottom tissue culture plates. Following the experimental treatments, cells underwent three washes with cold phosphate-buffered saline (PBS). Subsequently, cell viability was assessed using the Cell Count Reagent SF colorimetric assay (Nacalai Tesque, Kyoto, Japan). Briefly, 10 μL of Cell Count Reagent SF was added to each well, and the cells were incubated at 37°C for 2 hours. Cell viability was quantified colorimetrically by measuring OD450 values using a microplate reader (Model 550; Bio-Rad Laboratories, Hercules, CA, USA). The percentage of viable cells was calculated as follows: percentage viable cells = (T/C) × 100, where T and C represent the mean OD450 values of the treated and control groups, respectively.

### 2.8. Melanin Content Assay

To determine melanin content, cells were dissolved in 200 µL of 1 N NaOH for 30 min at 100^◦^C to solubilize the melanin, which was then quantified in cell suspensions by recording the absorbance at 405 nm as described previously^33^. Melanin content was calculated and corrected based on cell number.

### 2.9. Fluorescent immunohistochemical staining

Skin tissue samples were fixed in a 10% formaldehyde solution and subsequently embedded in paraffin. From these samples, 3-μm sections were prepared for fluorescent immunohistochemical staining. These sections underwent an overnight incubation at 4°C with primary antibodies specific to CYP27A1 ab126785, 1:100 dilution; Abcam) and Melan-A (M7196, 1:50 dilution; DAKO, Glostrup Kommune, Denmark). Subsequently, sections were treated with a secondary antibody (anti-rabbit IgG Alexa Fluor 488; anti-mouse IgG Alexa Fluor 555; Invitrogen, Thermo Fisher Scientific, Loughborough, United Kingdom). Additionally, sections were counterstained with Hoechst 33342 at a ratio of 1:500 (Invitrogen). The stained sections were visualized using either a light microscope or a Biozero 8100 confocal microscope (Keyence Co., Osaka, Japan).

### 2.10. Statistical analysis

Each experiment was replicated no less than three times. The data are displayed as mean ± standard deviation (SD). For the evaluation of interactions between variables, a two-way analysis of variance (ANOVA) was performed. To compare differences between two distinct groups, an unpaired Student’s *t*-test was applied using Microsoft Excel (Microsoft Corp., Redmond, WA, USA). Significance levels were set at *p*-values < 0.05, indicating statistical significance.

## 3. Results

### 3.1. Expression of core circadian genes BMAL1 and CLOCK in various human skin cell types

To determinate the peripheral expression of core circadian clock genes within human skin, we undertook a systematic investigation into the expression profiles of the key circadian clock components BMAL1 and CLOCK across fourteen distinct human skin cell types (Fig. 1H).

Quantitative real-time PCR revealed that both BMAL1 and CLOCK were expressed in all examined cell types, albeit at different levels (Fig. 1A, B). Among them, melanocytes exhibited markedly higher mRNA expression compared with fibroblasts or keratinocytes. Western blot analysis further confirmed the protein expression of BMAL1 and CLOCK across all skin cell types, with signal intensities largely consistent with the mRNA data (Fig. 1C, D, E). Immunofluorescence staining additionally demonstrated that BMAL1 and CLOCK were clearly detectable in melanocytes, showing predominant nuclear localization, consistent with their role as transcriptional regulators (Fig. 1F, G).

Collectively, these results indicate that the core clock components BMAL1 and CLOCK are expressed in multiple human skin cell types, with particularly high expression in melanocytes, suggesting a potentially important role of the circadian machinery in pigment cell physiology.

### 3.2. Efficient knockdown of BMAL1 and CLOCK in human melanocytes reduces melanin synthesis

To elucidate the functional role of circadian clock genes in melanocytes, siRNA-mediated knockdown of BMAL1 and CLOCK was performed in cultured human epidermal melanocytes (HEMn-MPs). Quantitative real-time PCR confirmed significant reductions in both BMAL1 and CLOCK transcript levels after siRNA transfection (Fig. 2A, B). Western blot analysis further showed marked decreases in BMAL1 and CLOCK protein expression in the knockdown groups (Fig. 2C).

**Figure 2.**
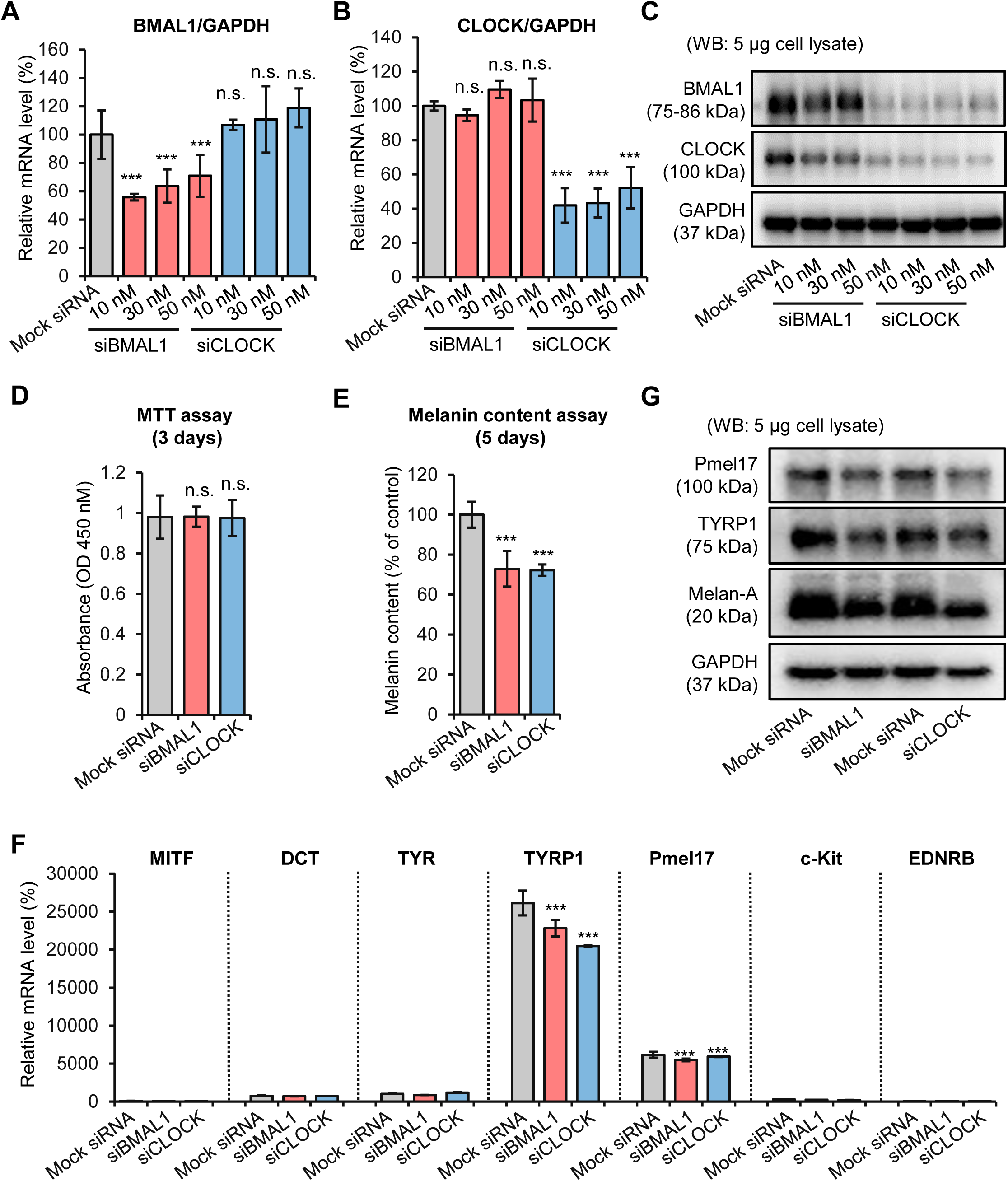
siRNA-mediated knockdown of BMAL1 and CLOCK reduces melanogenesis in human epidermal melanocytes (HEMn-MP). Quantitative real-time PCR analysis showing efficient suppression of BMAL1 **(A)** and CLOCK **(B)** mRNA expression 24 hours after transfection with siBMAL1 or siCLOCK (10, 30, or 50 nM). Gene expression levels were normalized to GAPDH. Western blot analysis of BMAL1 and CLOCK protein expression in HEMn-MP cells 48 hours after siRNA transfection. GAPDH served as the loading control **(C)**. Cell viability measured by MTT assay 72 hours after transfection, showing no significant cytotoxicity induced by BMAL1 or CLOCK knockdown **(D)**. Intracellular melanin content quantified 6 days after siRNA treatment, demonstrating reduced melanogenesis in BMAL1- and CLOCK-silenced melanocytes **(E)**. Quantitative real-time PCR analysis of key melanogenesis-related genes (MITF, DCT, TYR, TYRP1, Pmel17, c-Kit, and EDNRB) 24 hours after siRNA transfection **(F)**. Western blot validation of selected melanogenic proteins (Pmel17, TYRP1, Melan-A) showing decreased protein expression 48 hours following BMAL1 or CLOCK knockdown **(G)**. Data in **A**, **B**, **D**, **E**, and **F** are shown as mean ± SD. ***p < .01; n.s., no significant difference, as determined by Student’s t-test.

To determine whether silencing these clock genes affects melanocyte viability or proliferation, an MTT assay was conducted (Fig. 2D). No significant differences were observed among the mock-, siBMAL1-, and siCLOCK-transfected groups, indicating that the knockdown did not induce cytotoxicity or impair proliferation. Given the primary role of melanocytes in pigment production, we next examined the functional consequences of BMAL1 and CLOCK depletion on melanogenesis. Measurement of intracellular melanin content revealed a significant decrease in both siBMAL1-and siCLOCK-treated cells compared with mock-transfected controls (Fig. 2E).

To further examine the mechanisms underlying the reduced pigmentation, we analyzed the expression of key melanogenic genes. Quantitative real-time PCR revealed that the mRNA levels of major melanogenesis-associated genes, including TYR, TYRP1, DCT, and the melanosomal structural protein Pmel17, were markedly reduced following BMAL1 or CLOCK knockdown (Fig. 2F). Consistent with these transcriptional changes, western blot analysis demonstrated clear reductions in the protein levels of TYRP1, Pmel17, and Melan-A in both knockdown groups (Fig. 2G). These coordinated reductions indicate that BMAL1 or CLOCK suppression broadly attenuates the melanogenic pathway.

Together, these findings demonstrate that BMAL1 and CLOCK were effectively silenced in human melanocytes, and their downregulation led to reduced melanin synthesis through suppression of multiple melanogenic genes, without affecting cell viability. These results indicate that both BMAL1 and CLOCK act as positive regulators of melanogenic activity in human melanocytes.

### 3.3. Knockdown of BMAL1 and CLOCK selectively reduces CYP27A1 expression among multiple skin-related hormonal pathways

To explore whether the circadian clock influences local endocrine systems in melanocytes, we analyzed the expression of genes belonging to six major hormonal pathways relevant to skin physiology, including opsins, thyroid hormone signaling, melatonin synthesis, vitamin D3 metabolism, cortisol signaling, and estrogen signaling (Fig. 3A). Quantitative real-time PCR analysis revealed that the vast majority of genes across these pathways were unaffected by BMAL1 or CLOCK knockdown. However, a pronounced and selective decrease was observed in CYP27A1, a mitochondrial enzyme that catalyzes the initial hydroxylation of vitamin D3.

**Figure 3.**
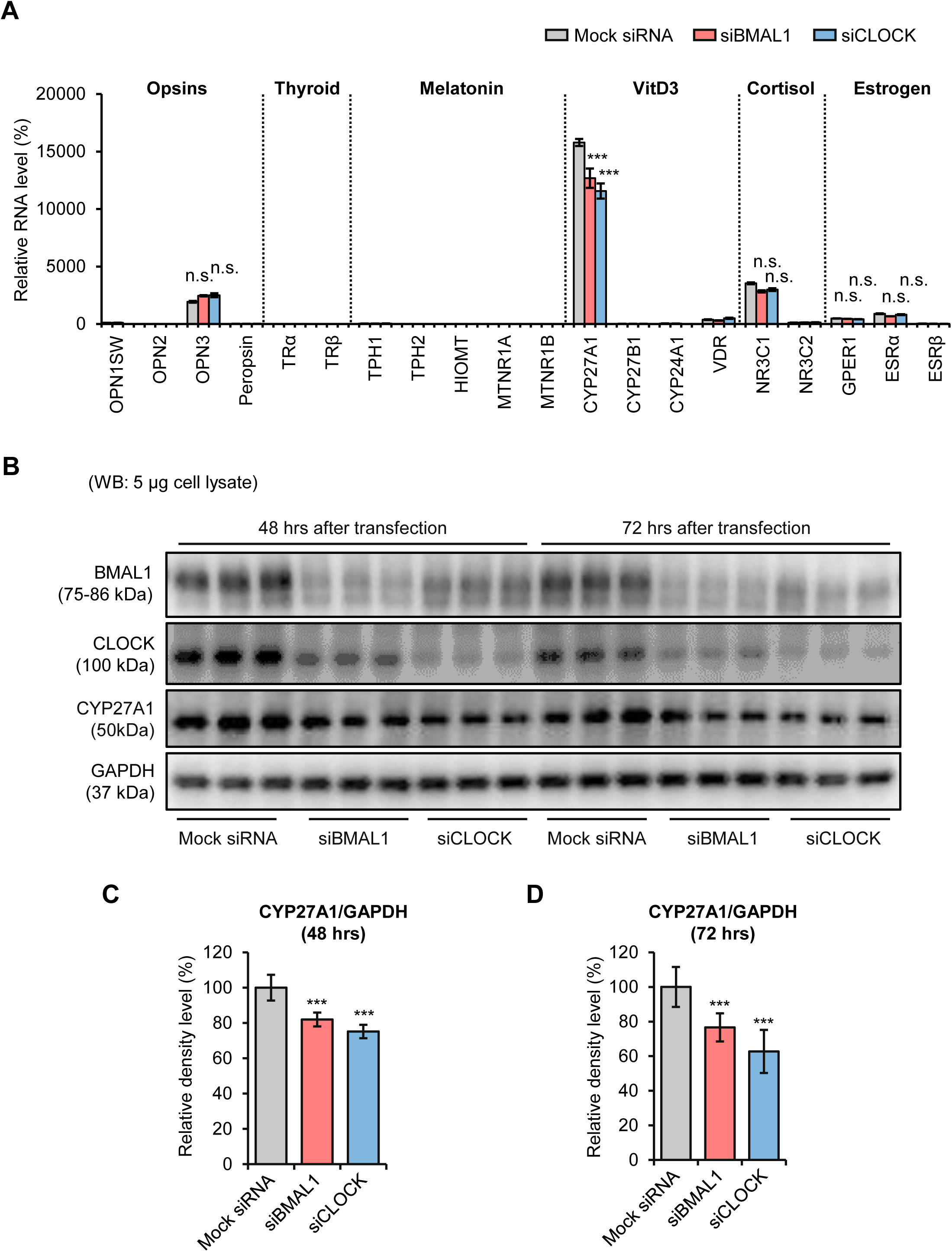
Knockdown of BMAL1 and CLOCK selectively suppresses CYP27A1 expression in human epidermal melanocytes. Quantitative real-time PCR analysis of hormone-related genes across six functional categories (opsins, thyroid signaling, melatonin pathway, vitamin D metabolism, cortisol signaling, and estrogen signaling) 24 hours after transfection with siBMAL1 or siCLOCK. Among all targets tested, CYP27A1 showed the most pronounced reduction following BMAL1 or CLOCK knockdown. Gene expression levels were normalized to GAPDH **(A)**. Western blot analysis of BMAL1, CLOCK, and CYP27A1 protein levels in HEMn-MP melanocytes at 48 and 72 hours after siRNA transfection. GAPDH served as the loading control **(B)**. Densitometric quantification of CYP27A1 protein normalized to GAPDH at 48 hours **(C)** and 72 hours **(D)** post-transfection, confirming a sustained reduction in CYP27A1 protein expression in BMAL1- and CLOCK-silenced cells. Data in **A**, **C**, and **D** are shown as mean ± SD. ***p < .01; n.s., no significant difference, as determined by Student’s t-test.

CYP27A1 converts vitamin D3 (cholecalciferol) to 25-hydroxyvitamin D3 [25(OH)D3], the principal circulating precursor of hormonally active calcitriol. Thus, the selective decrease in CYP27A1 suggests that circadian clock disruption may impair melanocyte-intrinsic vitamin D metabolic capacity.

To confirm the transcriptional results, CYP27A1 protein levels were examined by Western blotting at 48 and 72 hours after siRNA treatment (Fig. 3B). Consistent with the mRNA data, CYP27A1 protein expression was significantly reduced in both siBMAL1- and siCLOCK-treated melanocytes at both time points (Fig. 3C, D), while BMAL1 and CLOCK knockdown efficiency was validated by the corresponding protein depletion.

These results demonstrate that among multiple cutaneous hormonal pathways screened, CYP27A1 is uniquely sensitive to BMAL1/CLOCK depletion, indicating that the core circadian machinery may play a regulatory role in controlling melanocyte-intrinsic vitamin D3 hydroxylation capacity.

### 3.4. CYP27A1 expression is diminished in melanocytes within vitiligo lesions

To determine if the vitamin D3 metabolic pathway is dysregulated in vitiligo, we analyzed CYP27A1 protein expression in vitiligo patients and control epidermal tissues by immunofluorescence staining (Fig. 4). In healthy skin, robust CYP27A1 expression was observed in melanocytes, verified by co-localization with Melan-A. These CYP27A1-positive melanocytes were normally distributed along the basal layer. Compared to controls, vitiligo lesions displayed a marked decrease in Melan-A–positive melanocytes. Within the few remaining melanocytes, CYP27A1 signal intensity was noticeably reduced relative to control melanocytes.

**Figure 4.**
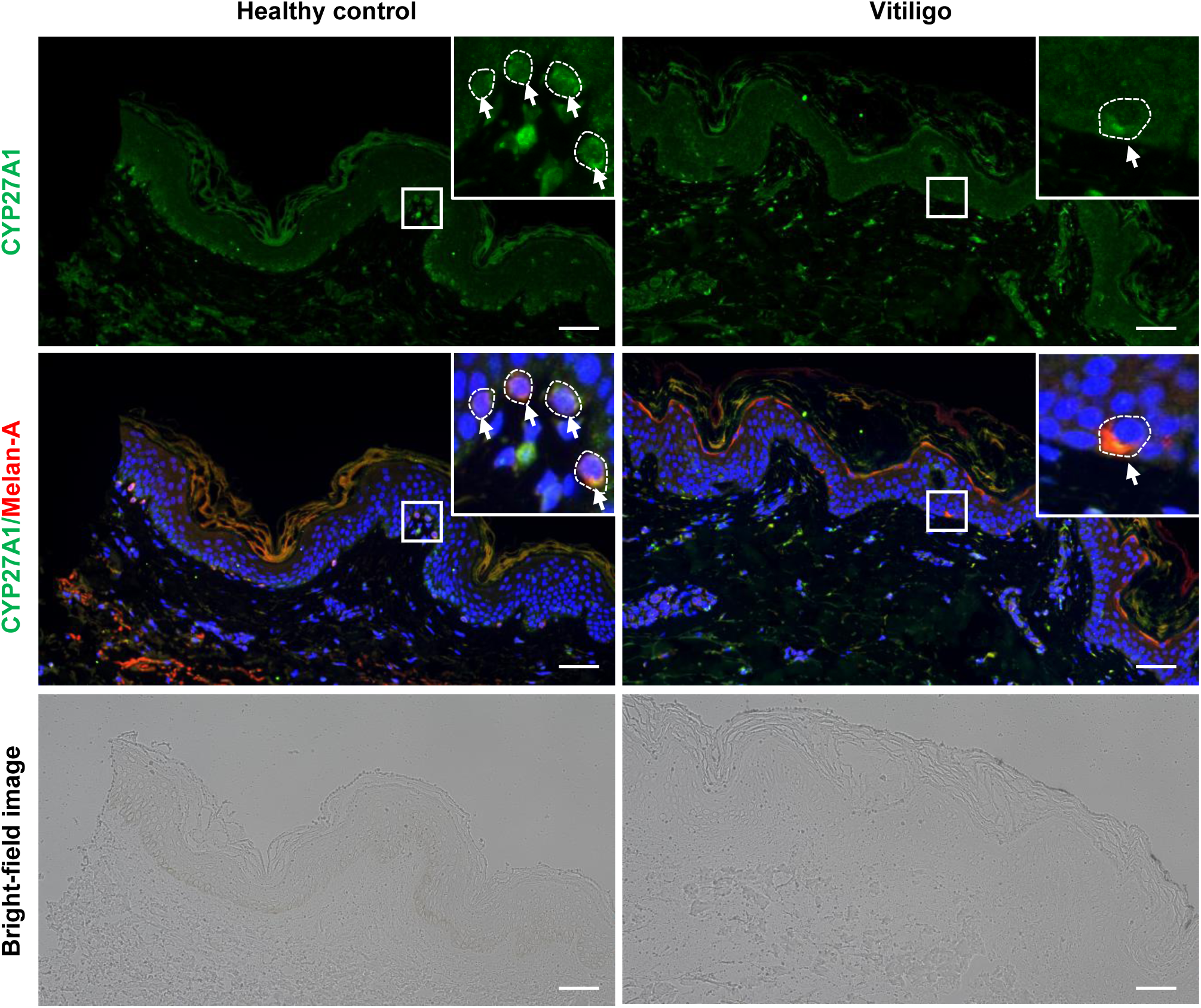
Immunofluorescence staining of CYP27A1 in vitiligo and healthy control skin. Representative immunofluorescence images showing CYP27A1 expression and Melan-A-positive melanocytes in lesional vitiligo skin compared with healthy control epidermis. CYP27A1 signals (green) and Melan-A (red) are shown individually and as merged images with DAPI (blue). Insets highlight melanocyte-existed areas in healthy skin and the reduced or absent melanocytes in vitiligo lesions. Corresponding bright-field images are provided for reference. Scale bars: 100 µm.

These observations indicate that CYP27A1 expression is diminished in the melanocytes of vitiligo lesions, correlating with the impaired vitamin D3 metabolism previously observed in BMAL1/CLOCK-knockdown melanocytes in vitro.

## 4. Discussion

Collectively, our integrated transcriptomic, proteomic, and spatial analyses provide compelling evidence that the core circadian regulators BMAL1 and CLOCK are predominantly and functionally expressed in epidermal melanocytes among the major human skin cell types examined. This expression profile strongly suggests that melanocytes possess an intrinsic and robust circadian clock, which may play a pivotal role in regulating local rhythmic functions in the skin.

The epidermis serves as the primary site for the initial photoconversion of 7-dehydrocholesterol to previtamin D₃ and subsequently to vitamin D₃ (cholecalciferol) upon UVB exposure^34^. Within epidermal cells, the mitochondrial enzyme CYP27A1 catalyzes the first critical step in vitamin D₃ activation, hydroxylating it to form 25-hydroxyvitamin D₃ [25(OH)D₃], the major circulating vitamin D metabolite and a key biomarker of vitamin D status^35^. Beyond its classical role in calcium homeostasis, locally synthesized vitamin D₃ exerts potent anti-inflammatory, immunomodulatory, and photoprotective effects, including the attenuation of sunburn cell formation and reduction of UV-induced cyclobutane pyrimidine dimers (CPDs), thereby contributing to cutaneous genomic stability and cancer prevention^36^.

Our functional studies demonstrate that BMAL1/CLOCK knockdown significantly suppresses CYP27A1 mRNA and protein expression in melanocytes, indicating that circadian disruption compromises melanocyte-dependent 25(OH)D₃ biosynthesis. These results reveal a previously unrecognized role of the circadian clock in regulating the local vitamin D metabolic axis within the epidermis. At the systemic level, impairment of this melanocyte-derived pathway may generate a dual vulnerability: weakened local immune and photoprotective defenses and reduced peripheral availability of 25(OH)D₃, the essential precursor for the active hormone 1,25(OH)₂D₃. Consistent with this, our immunohistochemical analysis demonstrates markedly reduced CYP27A1 expression in vitiligo lesions, providing new mechanistic insight that altered circadian regulation of vitamin D metabolism may contribute to the local microenvironmental dysfunction characteristic of depigmented skin.

Vitamin D₃ has long been reported to enhance melanocyte activity, promoting dendricity, increasing tyrosinase activity, and elevating melanin production in epidermal melanocytes^37^. In line with this, we observed reduced melanin synthesis following BMAL1/CLOCK knockdown in epidermal-derived melanocytes (HEMn-MP). However, this finding contrasts with observations in hair follicle melanocytes, where silencing BMAL1 or PER1 leads to enhanced tyrosinase activity, increased TYRP1/2 expression, and augmented pigmentation^32^. These divergent outcomes likely reflect the fundamentally different physiological roles of the two melanocyte populations: epidermal melanocytes govern constitutive pigmentation and UV-induced photoprotection, whereas hair follicle melanocytes primarily support cyclic hair pigmentation and operate within a distinct niche governed by the follicular stem cell environment.

Taken together, our findings underscore that the core circadian clock exerts lineage-specific, microenvironment-dependent control over melanocyte physiology, influencing both melanogenesis and local vitamin D metabolism. This dual regulatory function suggests that circadian dysregulation may contribute to pigmentary diseases such as vitiligo not only by altering melanogenic pathways but also by impairing local immunophysiological resilience via disrupted vitamin D biosynthesis. Future studies dissecting the cell-type-specific chromatin landscapes, signaling networks, and temporal dynamics that link clock components to melanocyte function will be essential for unraveling these context-dependent behaviors and for guiding the development of chronobiology-based therapeutic strategies for pigmentary disorders.

## Acknowledgements

We appreciate the cooperation and help provided by our secretary, Kumiko Mitsuyama.

